# TopoQual polishes circular consensus sequencing data and accurately predicts quality scores

**DOI:** 10.1101/2024.02.08.579541

**Authors:** Minindu Weerakoon, Sangjin Lee, Emily Mitchell, Haynes Heaton

## Abstract

**Summary:** Pacific Biosciences (PacBio) circular consensus sequencing (CCS) aka high fidelity (HiFi) technology has revolutionized modern genomics by producing long (10+kb) and highly accurate reads by sequencing circularized DNA molecules multiple times and combining them into a consensus sequence. Currently the accuracy and quality value estimation is more than sufficient for genome assembly and germline variant calling, but the estimated quality scores are not accurate enough for confident somatic variant calling on single reads. Here we introduce TopoQual, a tool utilizing partial order alignments (POA), topologically parallel bases, and deep learning to polish consensus sequences and more accurately predict base qualities. We correct ~31.9% of errors in PacBio consensus sequences and validate base qualities up to q59 which is one error in 0.9 million bases enabling accurate somatic variant calling with HiFi data.

**Availability and implementation:** The source code and installation instructions as well as validation dataset used are freely available at https://github.com/lorewar2/TopoQual

## 1 Introduction

Somatic variants, unlike germline variants, occur in a subset of cells. The fraction of cells in which a given somatic variant occurs affects our ability to sample it. And in order to be confident it is a true positive rather than an erroneous base, we often must sample it multiple times. In order to confidently call a somatic variant from a single DNA read, the error rate of that base must be significantly lower than the rate of somatic variants expected in that genome^1,2^.

In order to have high base accuracy, we must create a high signal to noise ratio (SNR) system. The size of a single nucleotide of DNA is smaller than the wavelength of light. This makes measuring the sequence of nucleotides of a single DNA molecule optically near the theoretical diffraction limit of detection^3,4^. To overcome this, historically each molecule was amplified by cloning, polymerase chain reaction (PCR)^5^, or bridge amplification^6^. This increases the SNR by measuring thousands or millions of nucleotides instead of a single one. However, amplification methods are not error-free and if an error occurs early on in these systems, the vast majority of molecules will have the erroneous nucleotide and we will confidently sequence this error^7,8^ putting a cap on the theoretical base accuracy.

In addition to this limitation, ever since Solexa introduced the Genome Analyzer in the early 2000s, the progression of DNA sequencing technologies has focused on data throughput over data quality. Data quality is, of course, multifactorial. For DNA sequencing it is a combination of the base level accuracy as well as the length of the read. In fact, the length of the sequence increases the theoretical information content exponentially while the base accuracy of each base does so only linearly. There are many repeats in genomes caused by a multitude of phenomena including but not limited to transposable elements^9^, a variety of duplication events^9^, and viral inserts^10^. To resolve a sequence, we must have reads greater than the length of these repeats in order to anchor on unique sequences^11^. In 2010, PacBio introduced their continuous long read (CLR) technology using sophisticated zero mode waveguides (ZMW) to limit the number of nucleotides in the detection space thus increasing the resolution very near the diffraction limit of light^3^. Over the next decade, they improved the processivity of their DNA polymerase enzyme as well as detector and software to create reads that were tens of kilobases long. This is in contrast to Illumina reads which were ~100-150 bases. The downside to this technology was that the bases had a high error rate (85-90% accurate)^12^. While the theoretical information content of these reads were very high due to their length, they were very challenging to work with computationally as they relied on extensive all-vs-all alignment and multiple alignment rather than fast exact-match seeding^11,13^.

More recently, Pacbio released their CCS/HiFi technology. In this technology, they attach a hairpin adapter to each end of the molecule creating a circular construct. Using strand-displacing DNA polymerase, they are able to sequence the full circular construct multiple times^4^. They then separate each subread of the forward and reverse strand and create a consensus sequence from them. This, along with improvements in DNA polymerase processivity allow for many passes on the same circular molecule. Because the errors on each sequencing pass on the molecule are largely independent of each other, the accuracy of the consensus sequence is largely only theoretically limited by the number of passes and our ability to create accurate multiple alignments of these subreads. This creates long (15kb+) and highly accurate (99.9+%) reads with some bases reaching a theoretical accuracy on the order of 1-10^-9 or higher assuming no upstream error sources, optimal multiple alignment, and truly independent measurements. This technology has revolutionized genome assembly^14^, structural variant analysis^15^, and other aspects of genomics. However, when Pacbio estimates the quality of each base, it gives many bases a Phred scaled quality of 93 (corresponding to an accuracy of 1-10^-9.5) and when compared to a sample with nearly perfect ground truth knowledge, these bases only validate at q45. Therefore, we cannot trust these quality score estimates. It is our goal here, to create a system which not only estimates these quality scores accurately, but also can correct some of the errors in the consensus base calling algorithms currently used. This will pave the way to allow for accurate somatic mutation calling with CCS data even when the somatic variant is only sampled by a single read in the sample.

## 2 Materials and Method

### 2.0 CCS library preparation and sequencing

Umbilical blood from a newborn female was collected in 40-60 mL lithium-heparin tubes, and processed for blood granulocyte isolation using Lymphoprep. High molecular weight (HMW) DNA was extracted from the granulocytes using the Qiagen MagAttract HMW DNA extraction kit (67563) and sheared into 16-20 kb DNA fragments using the Megaruptor 3 system (B06010003) with a speed setting of 30. CCS sequencing libraries were then prepared following the standard CCS library preparation protocol 1.0 (100-222-300), and sequenced on Sequel IIe instruments at the Wellcome Sanger Institute.

### 2.1 Overview

We present TopoQual, a tool for polishing the sequences and providing precise base quality scores through the utilization of parallel bases. The workflow of TopoQual is illustrated in Figure 1A. To begin, we perform POA multiple alignment of subreads with the current consensus. We then use our algorithm (topocut) to find the alternative, or parallel, bases of the calling base in the POA graph. The parallel bases, in conjunction with the trinucleotide sequence of the read and the target base’s quality score, are input to the deep learning model treating mismatch bases that are not a germline mutation as errors because the number of somatic mutations in our umbilical cord blood data is expected to be much smaller than the number of errors observed. The bases matching the reference genome which are also not part of a germline variant are treated as non-errors. This model then produces a corrected quality score. Additionally, the parallel bases from topocut are utilized to correct the original base call if an alternate base has a higher count than the original base. While the reference genome is necessary for the training of this model, it is not required for new datasets which can be corrected and base quality recalibrated with just the subreads.

**[Figure 1]:**
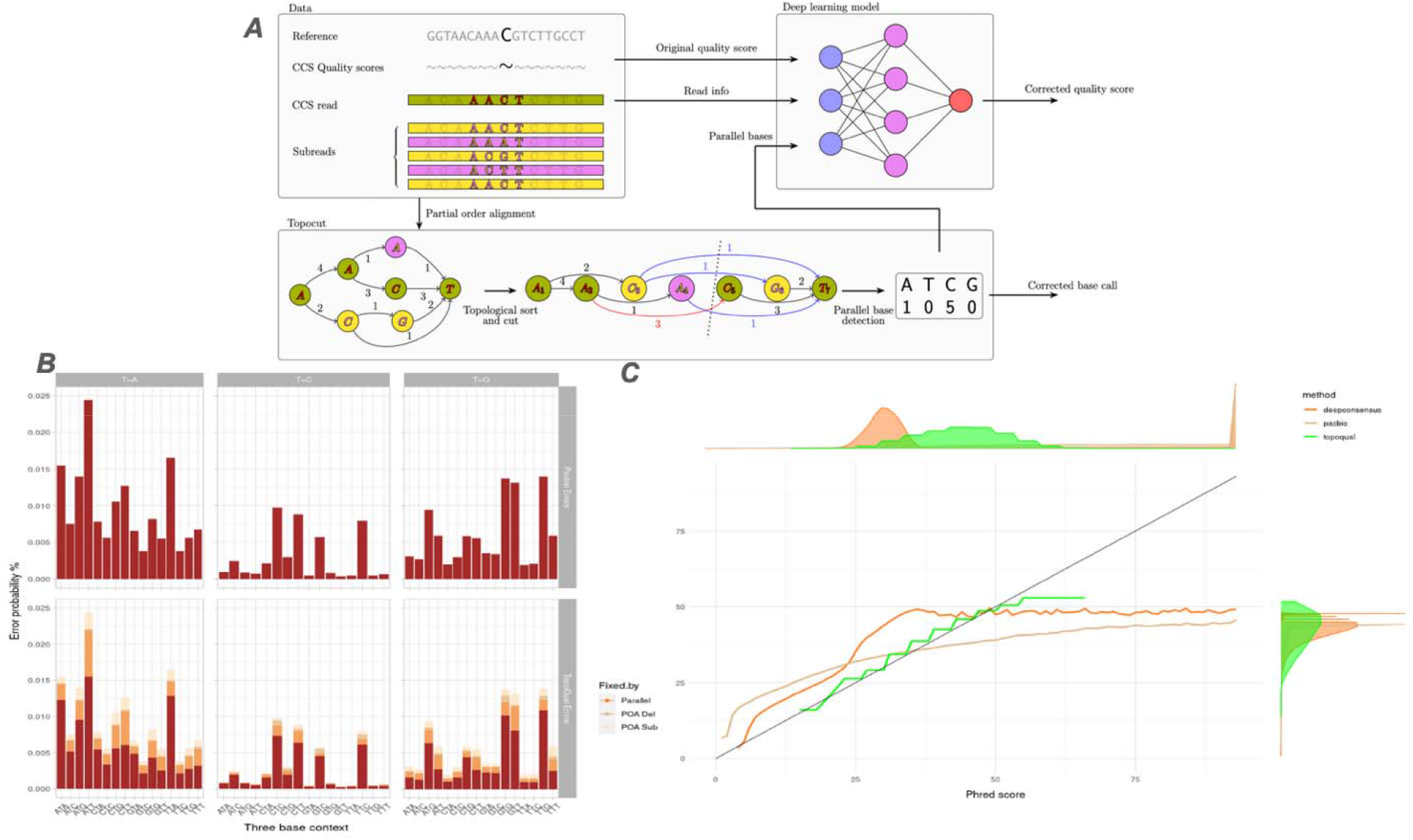
A. The quality score estimation strategy of topoqual. Subreads are aligned together with the current consensus sequence. Then potential alternative bases in the multiple alignment are detected via finding parallel bases in the POA graph (supp). These along with multiple other signals are sent to a deep learning system to learn a quality score estimator. B. Errors present before (top) and after polishing (bottom) by topoqual in the validation dataset T>X for our 3 types of corrections (parallel bases prefer a different base, POA consensus deletes a base, and POA consensus substitutes a base), chr2. C. Quality score validation comparison of different methods (expected to fall on the diagonal), chr2.

### 2.2 Topocut

The partial order alignment data structure^16^, which is a graph containing rich details about the aligned sequence structure, allows us to analyze the alternate pathways from the target base’s path; we define these alternate pathways as parallel bases. We use partial order alignment as it guarantees the optimal alignment of a new sequence versus the sequences already aligned. How Partial order alignment works is by extending standard dynamic programming sequence alignment^17,18^ to work with partial order graphs adding a sequence to the graph in each step.

TopoCut is the algorithm we used to procure parallel bases from the partial order alignment graph in our tool TopoQual. To accurately find the parallel bases, TopoCut first does partial order multiple sequence alignment with the CCS read and then the subreads. This outputs a partially ordered graph in which sequence letters are represented by nodes, and number of agreeing sequences are represented by edge weights. Then we sort nodes in a topological fashion and rank them according to the order. In this sorted graph, TopoCut makes a cut in front of the calling base and identifies the edges that intersect this cut, these edges are what we considered the parallel bases (figure 1A).

### 2.3 Deep learning model

The deep learning model is at the core of topoqual which takes in various information about the read and outputs the predicted quality score. Inputs to the model encompass the trinucleotide sequence of the read, Pacbio CCS quality score, parallel bases by topocut, average inter pulse duration, average pulse width, and the signal to noise ratio of the bases^19^. During the training phase, a dataset with labels of 0 for a reference mismatched base and 1 otherwise is utilized. Further details in the supplementary.

## 3 Results

### 3.1 Validation data

To validate our methodology, we sequenced a cord blood sample with few somatic mutations (40-50 somatic substitutions per cell^20^) from a 9-month-old female donor giving an expected number of somatic mutations in our 30x data of 675. Given the low mutation burden of this sample, most of the mismatches between the sample and the reference genome (524,575 observed) is a result of either library or sequencing errors, and not somatic mutations, indicating that the majority of these occurrences are likely attributed to erroneous base calls.

### 3.2 Sequence polishing

Topoqual conducts partial order alignment using PacBio CCS reads and their subreads to obtain parallel bases. Within this process, various mismatches (errors) with the reference are corrected using different techniques (parallel bases prefer a different base, POA consensus deletes a base, and POA consensus substitutes a base). Figure 1B illustrates the polishing of T>X mutations with respect to the three-base context. The percentages of errors corrected in different steps are as follows:

**[table1]:** Percentage of errors corrected by topoqual using different techniques, total number of bases analyzed 11Gb, errors present 1.8Mb.

| Polished by | Polished bases | Polished % |
| --- | --- | --- |
| POA SUB | 166Kb | 9 |
| POA DEL | 66Kb | 3.5 |
| PARALLEL | 361Kb | 19.5 |
| TOPOQUAL | 594Kb | 31.9 |

The sensitivity and specificity of sequence polishing are 31.9% and 99.6% respectively.

### 3.3 Comparison with Pacbio and Deep consensus quality scores

Pacbio and deep consensus uses Phred quality score outputs^421^ which range from 1 ~ 93, which corresponds to base call accuracy of 20% ~ 99.99999995%. Consensus and polishing algorithms seek to find the correct base as well as assign an accurate assessment of the likelihood of that base being erroneous. To do so, we count mismatches to the reference genome that are not germline variants as errors, but this method overlooks the presence of somatic mutations and considers them as errors. But because our validation dataset is from umbilical cord blood, the quantity of somatic mutations is much smaller than the number of observed mismatches (675 versus 524,575). This gives our validation a theoretical maximum quality value of q80 if the only mismatches we observed were somatic mutations (see supplement).

Figure 1C illustrates the algorithm-provided base quality scores (X-axis) compared to the corresponding actual base qualities from analyzing the mismatches in chromosome 2 of the validation dataset (Y-axis). The two marginal plots represent density distribution of the base counts.

At lower quality levels, both PacBio and DeepConsensus exhibit fewer errors than anticipated, but at higher quality levels, both surpass the expected error rates. PacBio reaches a maximum quality of 46, while DeepConsensus achieves 49. Our method, TopoQual, generates quality scores that align with the actual error numbers at both lower and higher quality levels, reaching a maximum of 54.

The distribution of quality scores in PacBio and DeepConsensus is predominantly concentrated around the maximum value, 93. However, the actual measured quality is well below 93. TopoQual more accurately measures the validated quality scores which are roughly normally distributed as expected. Despite a broader range of quality scores in TopoQual, the count of high-quality (>45) bases is equivalent to that of PacBio (±1%).

## 4 Conclusion

Correcting errors and providing accurate quality scores is necessary for single molecule sequencing somatic mutation calling. We introduce topoqual, a method for improving consensus sequence accuracy and dramatically increasing the validity of quality values. Topoqual corrects 31.9% of errors vs the PacBio consensus and produces accurate quality scores that have been validated versus a sample with exceedingly low somatic mutation burden. We show that existing methods highly overestimate the quality values of a majority of bases. Statistical methods overestimate base accuracy because of their assumption of total independence of subread sequences. Our quality values validate up to q59 or an error rate of 1e10^-5.9. This work will support the ability to accurately call somatic variants even when only one read samples the somatic variant.

