## Supplemental for "TopoQual polishes circular consensus sequencing data and accurately predicts quality scores"

**SUPPLEMENTARY**

**1. TOPOCUT Algorithm**

In our example (Main manuscript, Figure 1A), we aim to find the parallel bases of calling base C, which has a topological ranking of 5. First the parent edge weight of the calling base C is added to parallel base count. Then, parent-child rank pairs which sandwich the calling base C are discovered (3-6, 3-7, 4-7), and corresponding edge weights are added to the parallel base count to get the final parallel base count [A=1, C=5, G=0, T=0]. Total parallel base count is 7 which agrees with the total number of sequences, therefore further action is needed.

TOPOCUT_IDENTIFY_PARALLEL_BASES() (Figure1) accomplishes the above by, adding the calling base’s weight in graph to the parallel bases array (line 2) and adding the corresponding parent edge weights if there are any parent-child rank pairs that sandwich the calling base’s rank (line 3-7). If the count of parallel bases does not sum up to the total number of sequences Num (line 8 -9), the process is done in reverse (line 9-20).

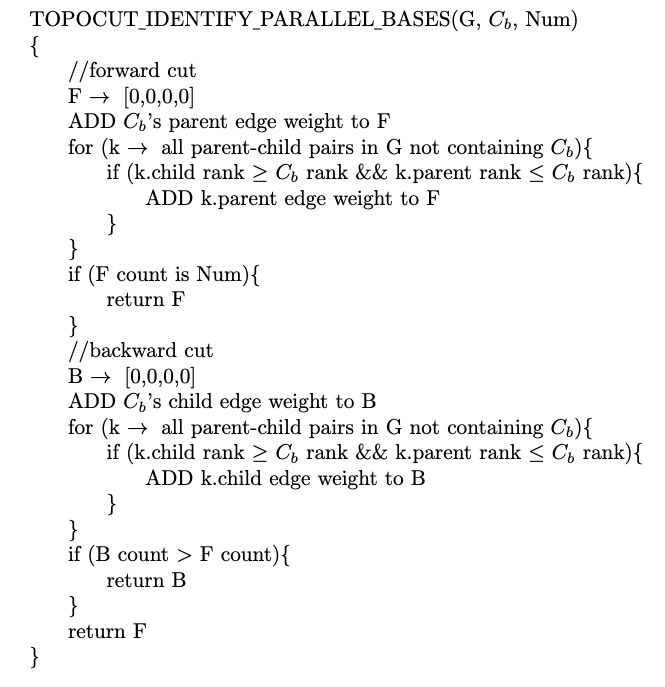

*[Figure 1]: Topocut algorithm pseudocode, G is the partial order graph, C_b_ is the calling base, and Num is the number of subreads for the read.*

**2. Bayesian estimation model**

**2.1 Quality calculation**

Bayesian estimation provides a method for incorporating prior knowledge into the estimation of unknown parameters^1^, in our case the unknown parameter is the error rate for the base-call. We denote the base-call as B_ij_, where i takes any value from the nucleotides A, C, G or T and j is the locus in the sequence. The prior probability, which is the probability of the base call, we set at p(B_ij_) = 0.25 for all locus. We use the probability model, binomial pdf, to get the probability of being correct given the base call,

$P(D|B_{i,j}) = \frac{n}{k}p^{k}{(1-p)}^{n - k}$ [1]

Where n and k are, respectively, the total number of sequences and number of sequences that agree with the base-call at locus j. Assumed accuracy of base-calling is denoted by p.

To get the correct rate for the base given the data we use,

$P(B_{i,j}|D) = \frac{P(D|B_{i,j})P(B_{i,j})}{\sum_{k\epsilon\{A,C,G,T\}} P(D|B_{k,j})P(B_{k,j})}$ [2]

This gives an estimation of the correct rate, this is converted to the standard of error probability measurement Phred score^2^.

${q_{j}=-10log}_{10}(1 -P(B_{i,j}|D))$ [3]

**2.2 Accuracy of base calling calculation.**

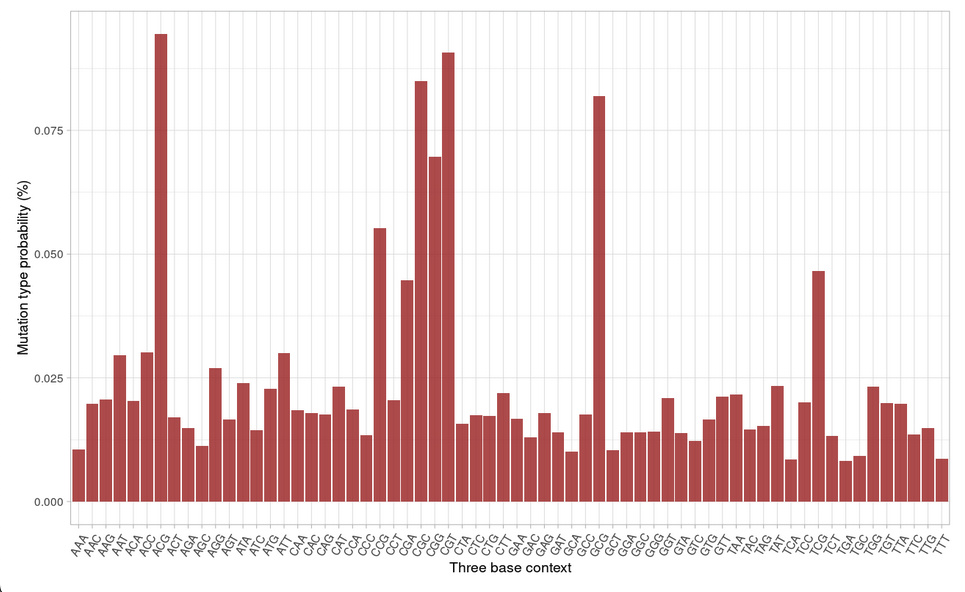

*[Figure 2]: Mutation type probability w.r.t three base context, chr1*

To obtain a variable accuracy of base calling **p**, we used the mutation count of three base contexts (Figure 2). First, we determined the average mutation probability for the dataset. Second, we obtained the three base contexts of the calling base and found the corresponding mutation type probability. Finally, we obtained the accuracy of base calling using the following formula:

${p=1 -(0.10)}(\frac{mutation type probability}{average mutation probability})$ [4]

Both the dynamic accuracy from [4] and a static accuracy of 0.85 were evaluated to determine the superior one.

**2.3 Results**

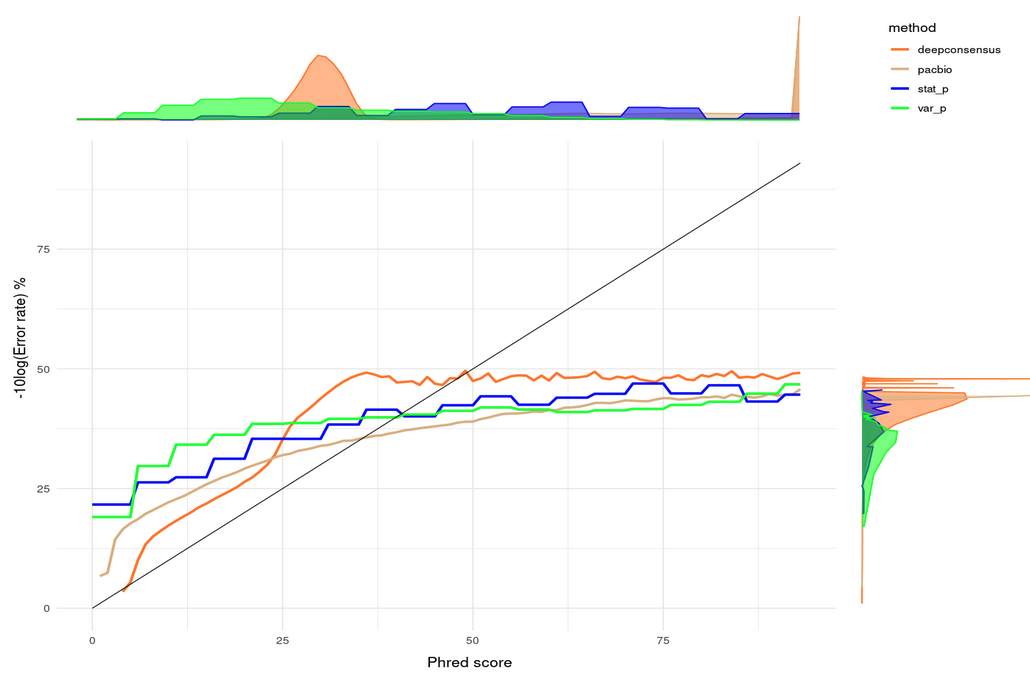

*[Figure 3]: Statistical model comparison graph, chr2*

As shown in Figure 3, using static ‘p’ yields higher counts of high-quality bases, but the distribution of bases is erratic. Conversely, using variable ‘p’ results in a lower count of high-quality bases, but they are normally distributed. These models, both static p and variable p, demonstrate fairly accurate predictions with an error probability of ~0.13%. However, they slightly underestimate base qualities up to 35 and then begin to overestimate them. This discrepancy is likely attributed to the strict independence assumption of this statistical model. For this reason, we developed a deep learning model to more accurately estimate base qualities.

**3. DEEP LEARNING MODEL DETAILS**

**3.1 DATA SET**

The acquired 9 month old cord blood granulocyte (PD47269d) with an average read length between 16kb and 20kb and coverage of 30x is our primary dataset, this sample only contains a few somatic mutations (281b / 90Gb) due to the average somatic mutation rate of 25 per cell per year^3^. Because we treat every mismatch that is not a germline mutation, our theoretical maximum quality value we could validate if we saw no mismatches that were not somatic variants would be q76.5.

Expected somatic mutations = 675b

Expected somatic mutations in the whole genome x 35% of the genome used for testing = 236b

Total bases used in testing = 10.67Gb

q80 = -10log10(236b/10.67Gb)

This dataset was split into different sets for the purpose of deep learning model creation.

| Set | Chromosome | Locus (Mb) |
| --- | --- | --- |
| Training set | Chr1 | 5 ~ 240 |
| Validation set | Chr1 | 240 ~ 250 |
| Test set | Chr2 | 5 ~ 240 |
|  | Chr3 | 5 ~ 200 |
|  | Chr4 | 5 ~ 190 |
|  | Chr18 | 5 ~ 80 |
|  | Chr19 | 5 ~ 58 |
|  | Chr20 | 5 ~ 64 |
|  | Chr21 | 5 ~ 45 |

*[table1]: dataset allocation*

**3.2 Preprocessing**

First, we need to filter out the germline variants in the dataset. If left unfiltered, these could be flagged as errors by our model. We use DeepVariant^4^, a state-of-the-art variant caller, to obtain a list of germline variant loci and then filter out those loci.

Second, we only consider bases that match with high-confidence regions of the GiaB reference. These regions are utilized in validating variant calling pipelines^5^, ensuring minimal errors in the reference.

Then, to filter out the trivial errors, we apply a number of hard filters to the error candidates (mismatched bases with reference). These filters are,

Trim filter: to filter out errors at the ends of the read, where read errors are common due to adapter trimming.

Indel filter: to filter out errors overlapping an indel site.

Low/High depth filter: to filter out the errors in low/high coverage regions, where reads could have been mapped incorrectly.

**3.3 Training**

The model and training parameters are configured as follows,

Optimizer : stochastic gradient descent

Loss function: mean squared error

Learning rate: 0.0001

Batch size: 1024 * 16

Epochs: 50

Fully connected layers: 3 (145 x 72 x 36)

**4. SUPPLEMENTARY FIGURES**

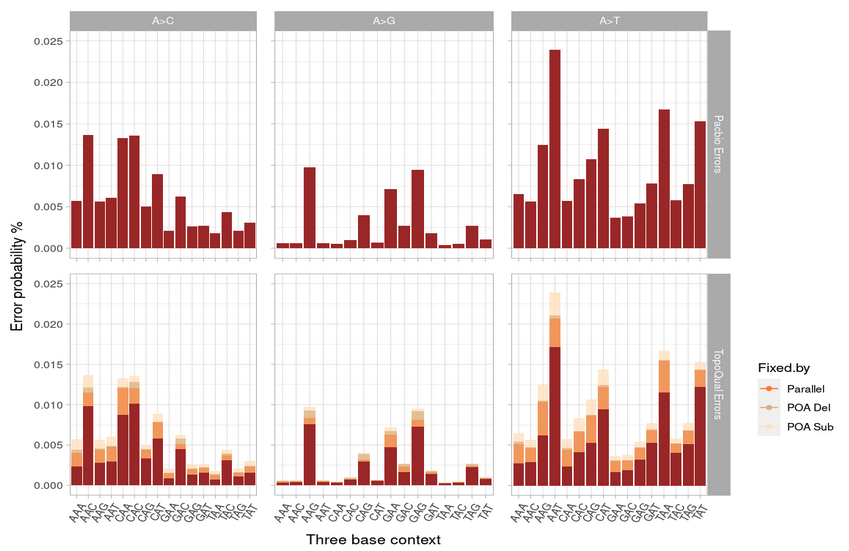

*[Figure 4]: Errors present before and after polishing by topoqual in the validation dataset A>X, chr2*

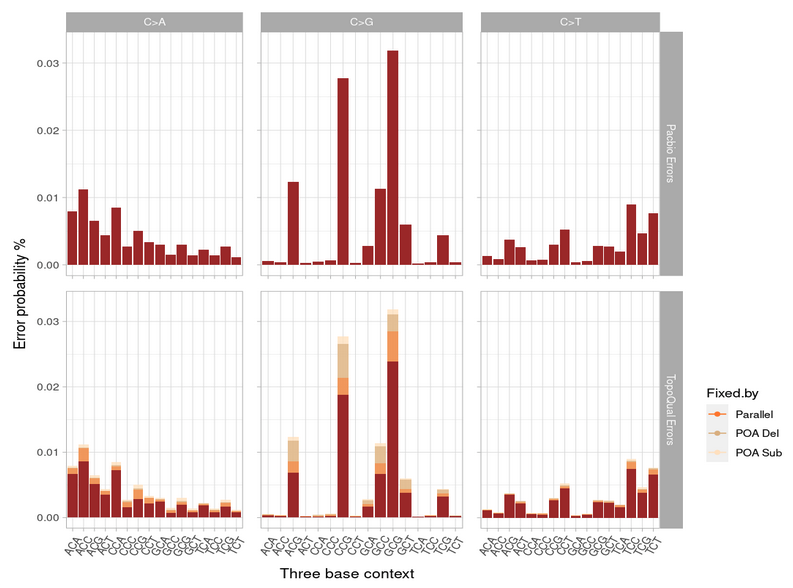

*[Figure 5]: Errors present before and after polishing by topoqual in the validation dataset C>X, chr2*

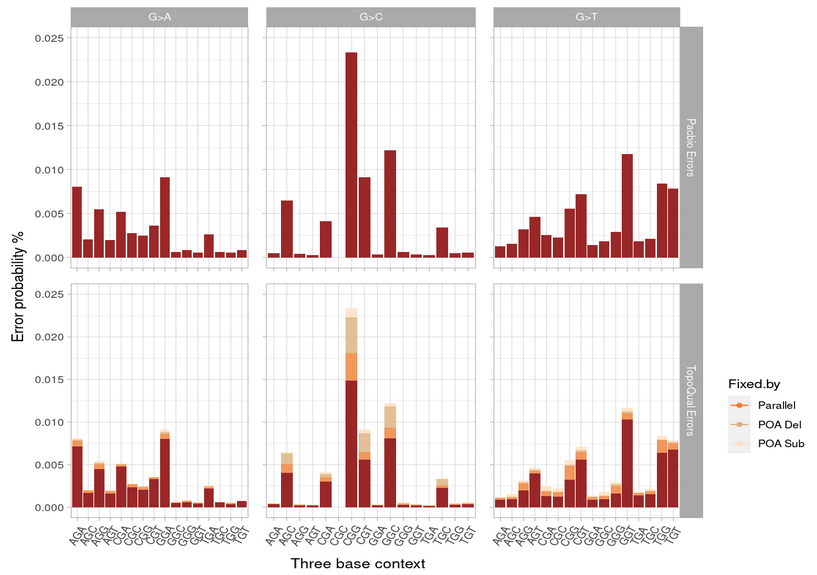

*[Figure 6]: Errors present before and after polishing by topoqual in the validation dataset G>X, chr2*

**5. SUPPLEMENTARY TABLES**

Average subread depth = 10

|  | **Pacbio** | | | **Deep Consensus** | | | **Topoqual** | | |
| --- | --- | --- | --- | --- | --- | --- | --- | --- | --- |
| **Chromosome** | **Error / Total base pairs** | **Error rate** | **Max Q** | **Error / Total base pairs** | **Error rate** | **Max Q** | **Error / Total base pairs** | **Error rate** | **Max Q** |
| Chr2 | 562Kb / 3.3Gb | 0.017% | 46 | 585Kb / 3.8Gb | 0.015% | 49 | 263Kb / 2.7Gb | 0.010% | 54 |
| Chr3 | 431Kb / 2.6Gb | 0.016% | 47 | 530Kb / 3.1Gb | 0.017% | 45 | 283Kb / 2.6Gb | 0.011% | 52 |
| Chr4 | 379Kb / 2.4Gb | 0.016% | 48 | 431Kb / 2.7Gb | 0.016% | 48 | 246Kb / 2.4Gb | 0.010% | 47 |
| Chr18 | 149Kb / 0.96Gb | 0.016% | 49 | 178Kb / 1.1Gb | 0.016% | 47 | 96Kb / 0.96Gb | 0.010% | 54 |
| Chr19 | 144Kb / 0.74Gb | 0.019% | 44 | 166Kb / 0.9Gb | 0.018% | 44 | 100Kb / 0.74Gb | 0.013% | 48 |
| Chr20 | 126Kb / 0.78Gb | 0.016% | 49 | 145Kb / 0.92Gb | 0.016% | 50 | 82Kb / 0.78Gb | 0.011% | 59 |
| Chr21 | 63Kb / 0.37Gb | 0.017% | 46 | 85Kb / 0.52Gb | 0.017% | 47 | 42Kb / 0.37Gb | 0.011% | 51 |

*[table2]: Results of Different methods on the validation dataset.*

**6. Code availability**

<https://github.com/lorewar2/TopoQual>
